# Coupling DNA processing to early gene expression drives antibiotic resistance plasmid dissemination

**DOI:** 10.64898/2026.01.09.698686

**Authors:** Nathan Fraikin, Yoshiharu Yamaichi, Christian Lesterlin

## Abstract

The rapid global spread of antibiotic resistance is mediated by conjugative plasmids, yet how these elements establish immediately after entering new bacterial hosts remains poorly understood. Here we show that plasmids actively coordinate DNA processing with early gene expression to promote their own establishment. Using single-cell measurements of plasmid conversion dynamics, we demonstrate that the F plasmid selectively delays complementary strand synthesis within its leading region, creating a transient single-stranded DNA window that prolongs zygotic induction. This delay is imposed by a single-stranded DNA promoter that forms a structural roadblock to complementary strand DNA synthesis and is resolved by the host helicase UvrD. By extending the single-stranded state of the leading region, plasmids enhance early expression of protective genes that counter host defence responses and promote survival during entry, particularly in non-isogenic hosts. Together, these findings identify early post-entry regulation as a critical determinant of plasmid establishment and reveal how coupling DNA conversion to transient protective gene expression enables conjugative plasmids to overcome host barriers and disseminate antibiotic resistance.

## INTRODUCTION

The global spread of antimicrobial resistance (AMR) represents a critical threat to public health and is primarily driven by horizontal gene transfer among bacteria (Munita Jose M. and Arias Cesar A., 2016). A major vehicle of this dissemination is bacterial conjugation, a contact-dependent process in which plasmids are transferred between cells (Castañeda-Barba et al., 2023). Surveys of resistant bacteria from environmental and clinical sources consistently show that the majority of acquired resistance genes are carried by conjugative plasmids, with plasmids of the IncF incompatibility group playing a disproportionate role in the spread of resistance and virulence traits among enterobacterial pathogens (Mathers Amy J. et al., 2015). Notably, it has been estimated that nearly two-thirds of acquired resistance genes in human-associated *Escherichia coli* reside on F-like plasmids, underscoring their epidemiological significance(Stephens et al., 2020) (Stephens et al., 2020).

Conjugative transfer proceeds through three sequential steps: formation of a mating pair, transfer of the plasmid as single-stranded DNA (ssDNA) into the recipient cell, and conversion of the incoming DNA into a double-stranded plasmid capable of replication and inheritance (Fraikin et al., 2024). While the molecular mechanisms governing plasmid processing and transfer in donor cells have been extensively studied, the molecular events that unfold within recipient cells immediately following plasmid entry remain far less well characterized. This early post-entry phase, termed plasmid establishment, has emerged as a critical and previously underappreciated bottleneck for plasmid dissemination (Fraikin et al., 2024). During establishment, plasmids enter recipient cells in a vulnerable ssDNA form and must rapidly counter host defence and stress responses while engaging host transcriptional and replication machineries to ensure stabilization. Accumulating evidence indicates that plasmids overcome this barrier through functions encoded within the leading region, the first plasmid segment to enter the recipient cell during conjugation (Fraikin et al., 2025b, 2024).

The leading region is uniquely positioned to act at the earliest stages of establishment and displays two defining features that enable the rapid deployment of protective functions (Frost et al., 1994). First, genes encoded in the leading region are expressed immediately upon plasmid entry, a phenomenon known as zygotic induction (Fraikin et al., 2025b; Jones et al., 1992). Zygotic induction was first described decades ago (Bates et al., 1999; Jones et al., 1992; Nasim et al., 2004), but its molecular basis has only recently been elucidated. We and others have shown that this early gene expression is driven by promoters that are active specifically on single-stranded DNA (Couturier et al., 2023; Masai and Arai, 1997; Wen et al., 2025). These ssDNA promoters adopt secondary structures that recruit RNA polymerase, allowing transcription to initiate as soon as the plasmid enters the recipient cell in its single-stranded form (Masai and Arai, 1997). Crucially, conversion of the plasmid into dsDNA inactivates these promoters, thereby terminating leading gene expression (Couturier et al., 2023; Masai and Arai, 1997). Thus, the duration of the zygotic program is intrinsically controlled by the timing of complementary strand synthesis. Second, the leading region encodes dense arrays of genes enriched in functions that promote plasmid survival during entry by counteracting host defence responses, including homologues of single-stranded DNA-binding proteins (Ssb), suppressors of the SOS response (PsiB), and multiple anti-defence systems such as inhibitors of restriction-modification systems, protective DNA methyltransferases, and inhibitors of CRISPR-based immunity (Samuel et al., 2024).

Together, these observations highlight a central paradox of plasmid establishment: incoming DNA must be converted to a double-stranded form to ensure stabilization and inheritance by its new host, yet the leading region must remain single-stranded long enough to sustain zygotic induction in a way that maximizes plasmid establishment success. Here, we address how plasmids resolve this antagonism by precisely tuning the spatiotemporal control of DNA conversion during conjugation, and how this balance critically determines establishment success.

## RESULTS

### Complementary DNA synthesis is selectively delayed in the F plasmid leading region

We first sought to quantitatively determine how rapidly different regions of the F plasmid are converted from single- to double-stranded DNA following entry into recipient cells. Plasmid transfer events were monitored using a translational fusion to the single-stranded DNA-binding protein Ssb in both donors and recipients, which form characteristic “kissing” foci on either side of the conjugation pore, as previously described (Couturier et al., 2023; Nolivos et al., 2019). Conversion to dsDNA was monitored by inserting *parS* sites at defined positions along the plasmid; recruitment of fluorescently labelled ParB proteins to these sites results in discrete foci upon dsDNA formation. Thus, the time delay between the appearance of Ssb and ParB foci provides a direct single-cell measurement of ssDNA-to-dsDNA conversion kinetics at each locus.

We first examined conversion kinetics in the maintenance, variable, and transfer regions by inserting four *parS* sites: one in the maintenance region, one in the variable region, and two in the transfer region (Figure 1a). Conversion of these four loci occurred with timings proportional to their distance from the origin of transfer (*oriT*) (Figure 1b). Linear regression of conversion delay as a function of distance from *oriT* yielded a conversion rate of 791 nucleotides per second, closely matching the historical estimate of 770 nucleotides per second derived from Hfr mating experiments (Wollman and Jacobs, 1958). These results indicate that, outside the leading region, complementary strand synthesis proceeds continuously and concomitantly with plasmid entry.

**Fig. 1:**
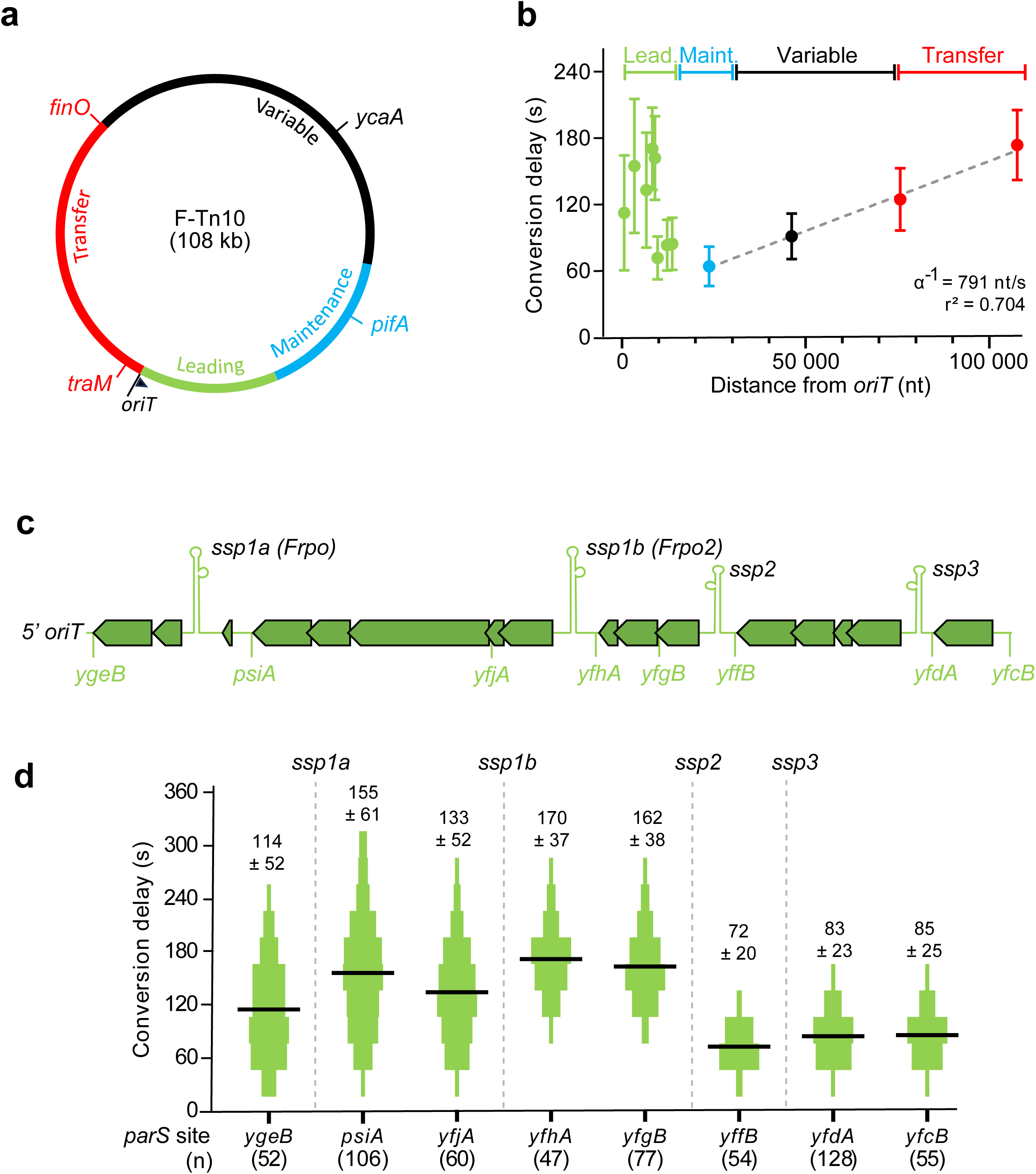
Conversion timing of the F plasmid during conjugation. **a** Genetic map of the F-Tn*10* plasmid indicating the leading (green), maintenance (blue), variable (black) and transfer (red) regions, as well as the origin of transfer (*oriT*), the direction of transfer (black arrow) and the position of *parS* insertions. **b** Conversion delay between the appearance of Ssb-mLychee and ParB-mChartreuse foci in transconjugant cells after the acquisition of the F-Tn10 plasmid labelled with *parS* sites at given coordinates in the leading (lead.), maintenance (maint.), variable or transfer region. Data shows the mean delay with standard deviations as error bars. A linear regression of conversion events in the maintenance, variable and transfer region is shown as a dashed lined with transfer speed estimated from the inverse of the regression slope (α). **c** Genetic map of the leading region showing open reading frames (arrows), single-stranded promoters and *parS* insertions. **d** Detailed conversion delays for *parS* sites inserted in the leading region. Data is shown as histograms of conversion delays measured for 3 independent experiments, with bin size representing the 30 s delay between image acquisitions. Mean conversion delays are shown as black bars and sample sizes are shown in brackets. Numbers show mean conversion times ± standard deviation.

We next focused on the leading region. Based on our recent work showing that the leading region is subdivided into four operons, each transcribed from a single-stranded DNA promoter, we inserted at least one *parS* site within each operon (Figure 1c). Strikingly, all *parS* sites located within the leading region were converted substantially later than predicted by their distance from *oriT*, revealing a clear deviation from the linear conversion kinetics observed elsewhere on the plasmid (Figure 1b). Moreover, conversion followed two distinct temporal regimes, with oriT-proximal loci converting markedly later (120-180 s) than oriT-distal loci (60-90 s) (Figure 1d). This pattern indicates that complementary strand synthesis is selectively and actively delayed within the leading region. Notably, we observed an abrupt, approximately two-fold increase in conversion delay between a *parS* site located adjacent to *yffB* (72 ± 20 s) and a downstream site adjacent to *yfgB* (162 ± 38 s). These two sites are separated by only 770 nucleotides but flank *ssp2*, a recently identified single-stranded DNA promoter (Wen et al., 2025). This sharp discontinuity suggested the presence of a localized cis-acting barrier to complementary strand synthesis at or near *ssp2*.

### A single-stranded DNA promoter acts as a replication roadblock during plasmid conversion

To test whether *ssp2* directly impedes progression of complementary strand synthesis, we sought to map the 3′ end of stalled DNA synthesis during conjugative transfer. DNA was extracted from mating cells, and putative stalled 3′ ends were extended *in vitro* with cytidine residues using terminyl transferase (Figure 2a). These extended products were then amplified by PCR using a poly[d(G-I)] primer in combination with a locus-specific primer (Figure 2a). A discrete 3′ end was captured from matings involving a wild-type F-Tn10 plasmid but not from matings with a conjugation-defective *traI*-deleted F-Tn10 plasmid, demonstrating that replication stalling occurs specifically during ssDNA-to-dsDNA conversion and not during vegetative plasmid replication (Figure 2b). Sanger sequencing mapped the stalled 3′ end precisely to the end of the predicted stem region of *ssp2*, implicating the stem-loop structure formed by this ssDNA promoter as a physical barrier to complementary DNA synthesis (Figure 2c abd 3d).

**Fig. 2:**
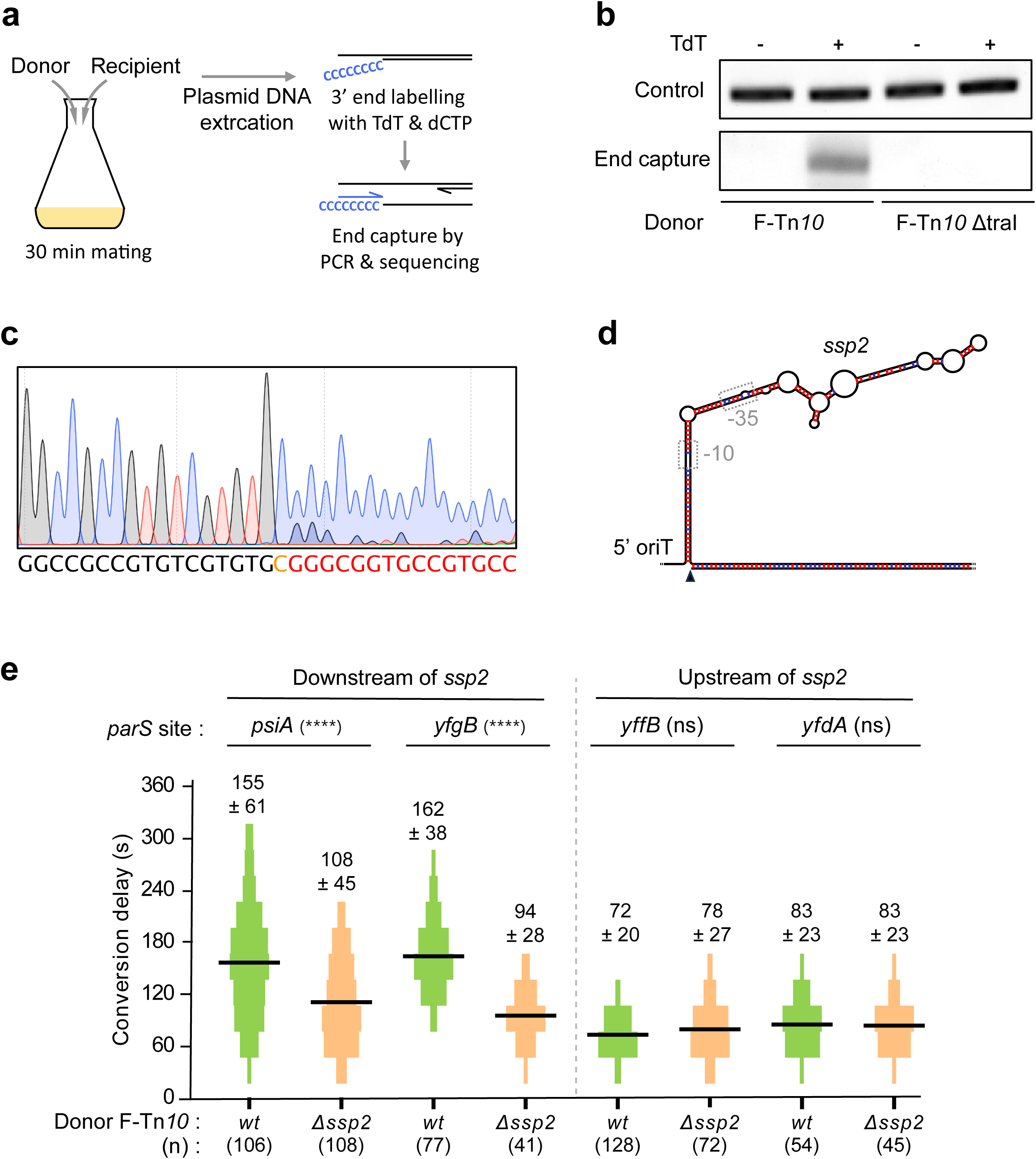
Identification of *ssp2* as a roadblock for F plasmid conversion. **a** Schematic representation of the setup used to map the 3’ end of a putative conversion intermediate. After a 30-min mating, plasmid DNA is extracted and 3’ ends are elongated with poly-d[C] using terminyl transferase (TdT) and dCTP. The 3’ end is captured by PCR using a poly-d[GI] anchor primer coupled to a primer that hybridizes to the locus of interest. After a second nested PCR, the amplicon is purified and sequenced. **b** Amplification products of the 3’ end capture experiments performed without (-) or with (+) terminyl transferase (TdT). Top row shows a control PCR. Bottom row shows the end capture with a poly-d[GI] anchor primer. **c** Sequencing trace analysis of the end capture product. Alignment with plasmid sequence is shown below. **d** Position of the mapped 3’ end on the putative secondary structure of *ssp2* as predicted by RNAfold (Lorenz et al., 2011). Red bars show G:C pairs, blue bars show A:T pairs and dashed boxes show the position of promoter elements. **e** Effect of a *ssp2* deletion on conversion timing in the leading region. Studied *parS* insertions are shown on top. Dashed line denotes the position of *ssp2*. Data is shown as histograms of conversion delays measured for 3 independent experiments, with bin size representing the 30 s delay between image acquisitions. Mean conversion delays are shown as black bars and sample sizes are shown in brackets. Numbers show mean conversion times ± standard deviation. ****: p<0.0001, Kruskal-Wallis with Dunn’s multiple comparison test, ns: p>0.05. Data for wt samples has been recapitulated from Fig. 1.

To directly test whether *ssp2* delays conversion, we deleted *ssp2* and measured conversion times at loci upstream and downstream of the promoter. Conversion times at *parS* sites upstream of *ssp2* (adjacent to *yfdA* and *yffB*) were unaffected by *ssp2* deletion (Figure 2d). In contrast, conversion of downstream loci (*yfgB* and *psiA*) occurred at significantly faster rate in the absence of *ssp2* (94 s vs 162 s for *yfgB*, 108 s vs 155 s for *psiA*). These results demonstrate that *ssp2* functions as a *cis*-acting replication roadblock that selectively delays complementary strand synthesis within the leading region.

### Transcription contributes to complementary strand synthesis but not to the conversion delay

Because the conversion delay observed in the leading region coincides with transcriptionally active stem-loop structures, we next asked whether transcription itself contributes to the observed delay in complementary strand synthesis. To address this question, we examined conversion kinetics in the presence of rifampicin, which inhibits RNA polymerase activity (Figure 3a). Rifampicin treatment did not abolish complementary strand synthesis, demonstrating that *de novo* synthesis of plasmid-encoded proteins is not required for plasmid conversion. This observation is consistent with earlier work showing that complementary strand synthesis in recipient cells relies exclusively on host-encoded replication factors (Hiraga and Saitoh, 1975). Importantly, rifampicin treatment did not suppress the conversion delay at *ssp2*, ruling out replication-transcription conflicts as well as plasmid-encoded factors as the cause of stalled DNA synthesis at this promoter.

**Fig. 3:**
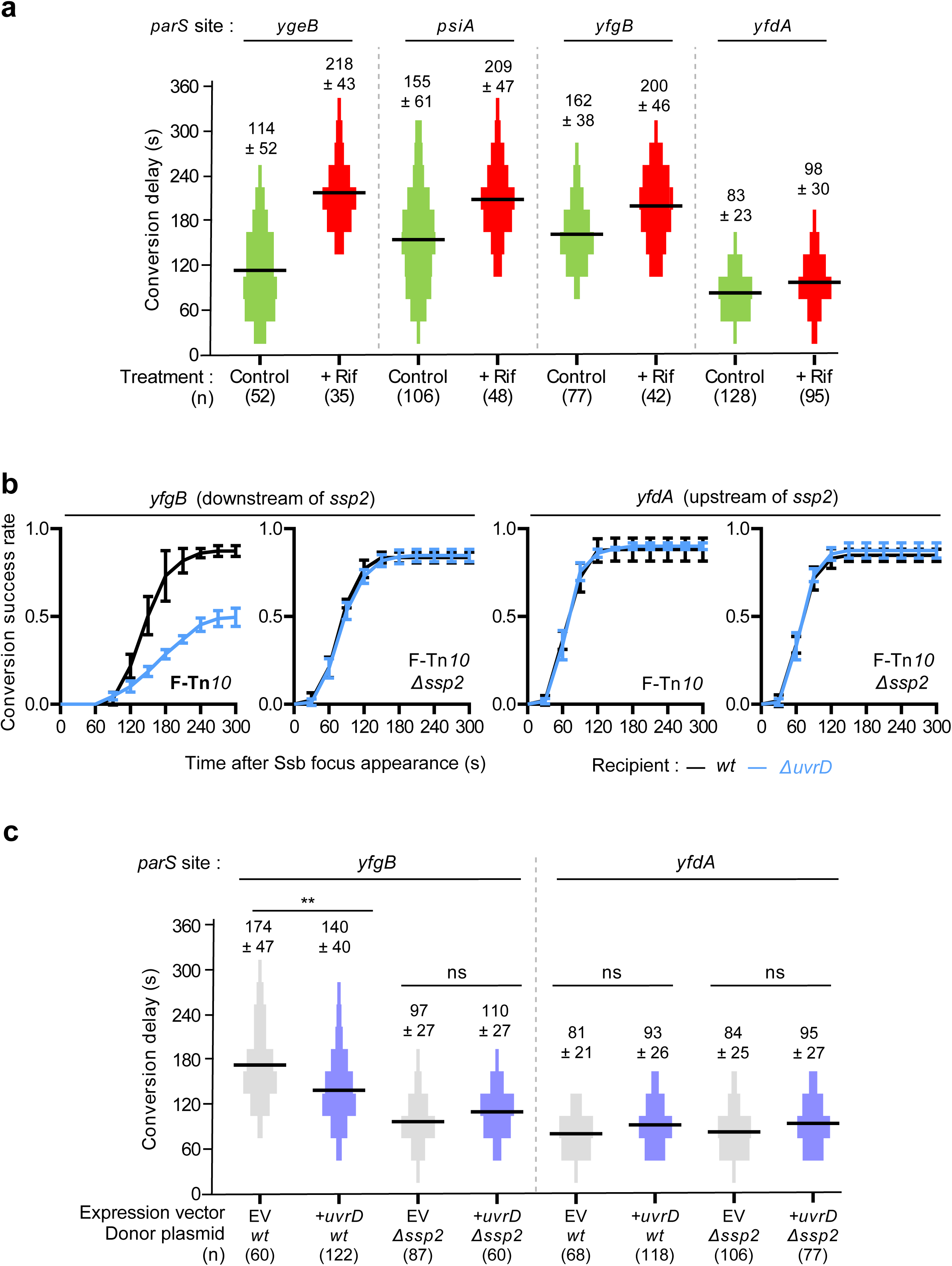
Effect of transcription inhibition and *uvrD* deletion on the *ssp2* roadblock. **a** Conversion delays for selected *parS* sites inserted in the leading region in the absence (green) or presence (red) of 200 µg/ml rifampicin. Data in absence of rifampicin is recapitulated from panel 1d. **b** Effect of a *uvrD* deletion on the conversion success rate of F-Tn*10*. Data shows the mean & SD of cumulative appearance delay between Ssb-mLychee foci and ParB-mChartreuse foci in wild-type (black) and Δ*uvrD* (green) recipients for three independent experiments. **c** Effect of *uvrD* overexpression on conversion timing in the leading region. Data is shown as histograms of conversion delays measured in at least 3 independent experiments, with bin size representing the 30 s delay between image acquisitions. Mean conversion delays are shown as black bars and sample sizes are shown in brackets. Numbers show mean conversion times ± standard deviation. ****: p<0.0001, ns p>0.05; Kruskal-Wallis with Dunn’s multiple comparison test.

In the absence of rifampicin, the distribution of conversion timings for loci located downstream of *ssp1b* (*psiA*) and *ssp1a* (*ygeB*) was markedly skewed towards fast conversion times compared to a *parS* site located upstream of ssp1b (*yfgB*) (Figure 1d & 3a). About 4% of conversion events at *yfgB* occurred at or before 90s versus 24% and 48% of events at *psiA* and *ygeB*, respectively, suggesting that conversion can be initiated at these loci (Figure 3a). Addition of rifampicin abolishes these early conversion events, suggesting that they are transcription-dependent. These findings are consistent with previous reports showing that transcription from *ssp1*/F*rpo* promoters can generate RNA-DNA hybrids that can act as primers for complementary strand synthesis *in vitro*. Together, these results provide an *in vivo* validation that transcripts from *ssp1*/F*rpo* promoters can be used as primers for complementary strand synthesis, while the conversion delay at *ssp2* arises from a structural replication barrier rather than transcription-replication interference.

### The host helicase UvrD enables progression through the leading-region barrier

Because stalling at *ssp2* is transient and resolves within 60-90 s, we hypothesized that a host-encoded activity actively dismantles the stem-loop structure that impedes DNA synthesis and thereby permits replisome progression. Among potential candidate factors, we focused on the host helicase UvrD, which has been previously implicated in the establishment of conjugative plasmids (Shen et al., 2023) and is well known for its ability to unwind DNA secondary structures and facilitate replisome progression across replication barriers (Bidnenko et al., 2006; Florés et al., 2005).

To test whether UvrD is required for conversion through *ssp2*, we deleted *uvrD* and measured both conversion success, defined as the fraction of ssDNA entry events (Ssb foci formation) that lead to dsDNA plasmid (ParB foci formation) (Figure 3b). In wild-type recipients, conversion success was 85% at both upstream (*yfdA*) and downstream (*yfgB*) loci. In Δ*uvrD* recipients, conversion upstream of *ssp2* was unaffected, whereas conversion downstream of *ssp2* exhibited both a reduced success rate (decreasing from 87% in wild-type cells to 49% in Δ*uvrD* recipients) and a significant delay among successful events (184 s versus 162 s), demonstrating that UvrD is specifically required to overcome the *ssp2*-dependent stall (Figure 3b). Consistently, deletion of *ssp2* restored both conversion success and normal conversion timing in Δ*uvrD* recipients (85% and 99 s, respectively).

We next asked whether increased UvrD availability could modulate progression through *ssp2*. Overexpression of *uvrD* from a multicopy plasmid selectively accelerated conversion downstream of *ssp2* (*yfgB*), reducing the mean conversion delay from 174 s to 140 s, whereas conversion upstream (*yfdA*) remained unchanged (Figure 3c). Importantly, this acceleration was lost when *ssp2* was deleted, confirming that UvrD acts specifically to resolve the *ssp2*-mediated roadblock rather than globally accelerating DNA synthesis. Together, these data identify UvrD as a host factor that enables efficient plasmid conversion by resolving a self-imposed structural barrier within the leading region.

### Delayed DNA conversion extends zygotic induction from single-stranded promoters

To assess how delayed conversion influences early gene expression, we fused the *psiB* gene, controlled by the *ssp1b*/F*rpo2* single-stranded promoter, to the fluorescent reporter mChartreuse (Fraikin et al., 2025a). Fluorescence was quantified in individual transconjugant cells at a defined time point (20 min) after detection of Ssb-mLychee foci (Figure 4a). Transconjugants receiving a wild-type F plasmid displayed significantly higher PsiB-mChartreuse fluorescence than those receiving an *ssp2*-deleted plasmid, with fluorescence intensities of 197 ± 142 a.u. and 101 ± 105 a.u, respectively (Pvalue <0.0001). This corresponds to an approximate 2-fold reduction in PsiB production upon *ssp2* deletion. These results indicate that delayed conversion of the leading region prolongs the single-stranded state of downstream loci, thereby extending the duration and magnitude of transcription from ssDNA promoters and enhancing early gene expression.

**Fig. 4:**
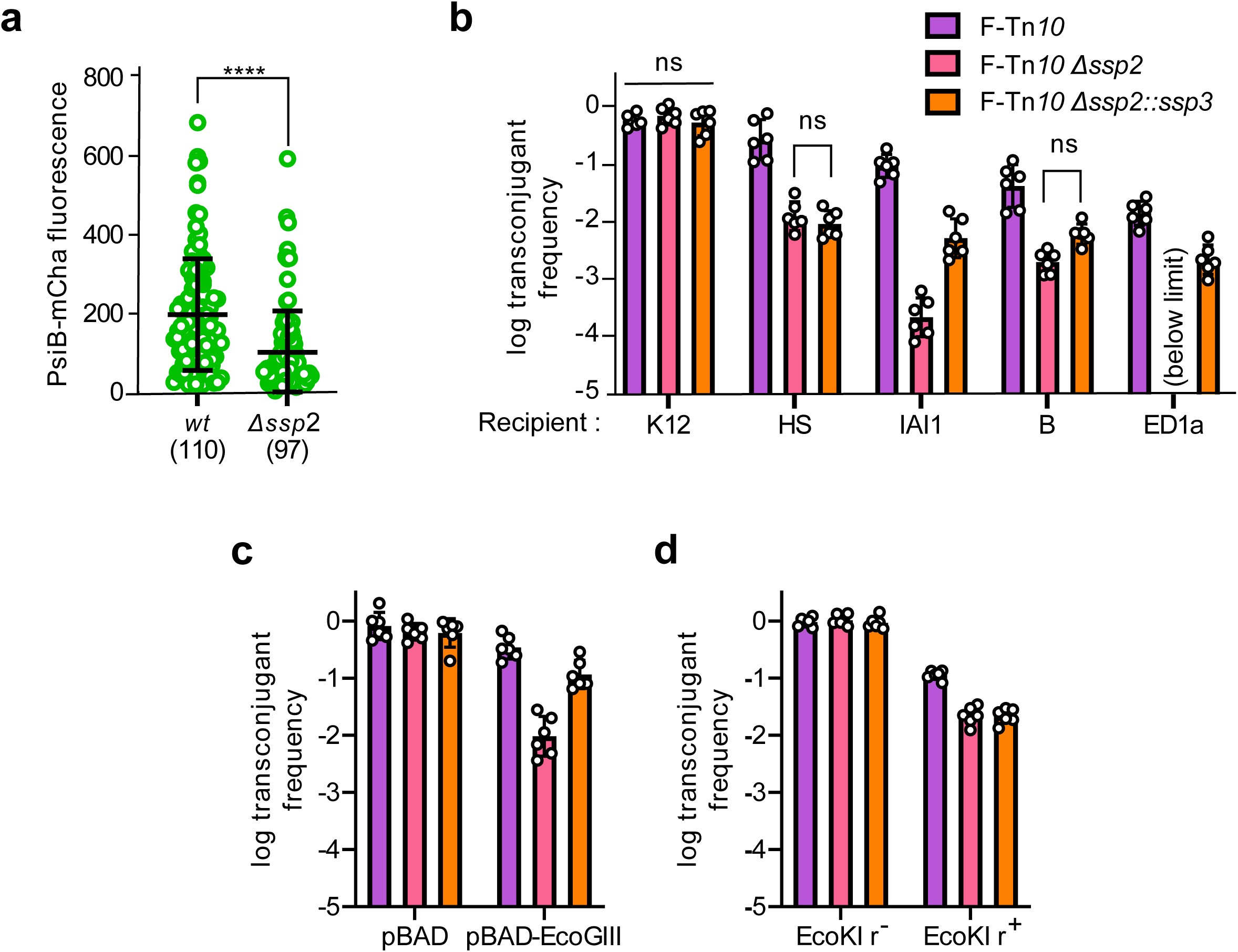
Effect of *ssp2* roadblock abolition on F-Tn*10* transfer efficiency. **a** Effect of *ssp2* deletion on the production of PsiB-mChartreuse. Data shows the fluorescence signal of single recipient bacteria 20 min after the appearance of Ssb-mLychee foci following the acquisition of a *psiB-mChartreuse* F plasmid. Bars show the mean and standard deviation of fluorescent signals. **** : P < 0.0001, Mann-Whitney test. **b** Effect of *ssp2* deletion and replacement with a *ssp3* copy on transconjugant frequency for transfer from *E. coli* K12 to the indicated *E. coli* strains on solid medium. Bars show the mean and SD of six independent experiments. ns : non-significant; Brown-Forsythe ANOVA with Dunnett’s T3 multiple comparison test. **c-d** Effect of *ssp2* deletion and replacement with a *ssp3* copy on transconjugant frequency for transfer of a plasmid to cells encoding the EcoGIII type II RM systems (**c**) or the EcoKI type I RM system (**d**). Bars show the mean and SD of six independent experiments. ns : non-significant; Brown-Forsythe ANOVA with Dunnett’s T3 multiple comparison test.

**Fig. 5:**
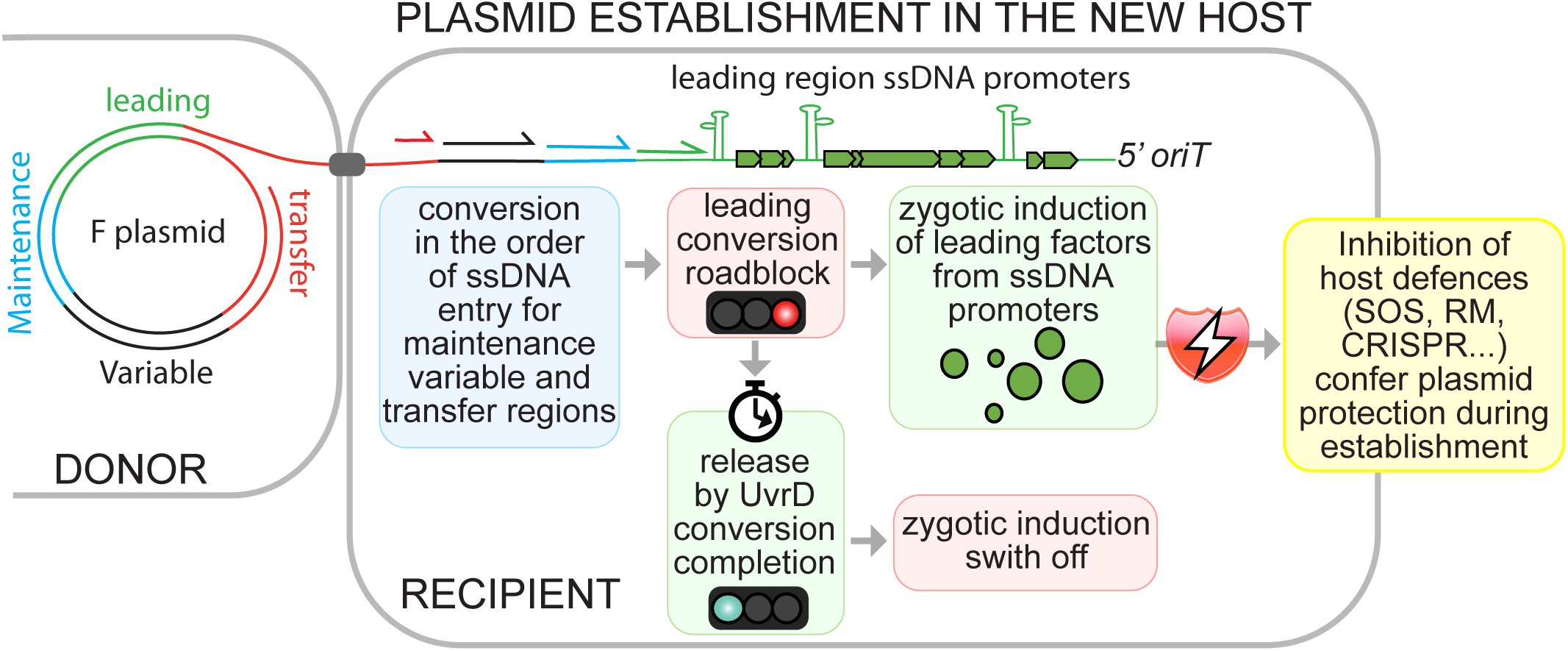
Working model. Conversion from ss-to-dsDNA operates in a discontinuous manner and hierarchically follows plasmid entry order for loci located outside of the leading region, *i.e.* the maintenance, variable and transfer. Conversion is stalled at the *ssp2* single-stranded promoter due to structural constraints, which increases residency time for ssDNA promoters, enables higher levels of zygotic induction, facilitates the inhibition of host defences and increases plasmid establishment success rate. Recruitment of the UvrD host helicase enables the melting of the *ssp2* stem-loop, facilitating replisome progression through this structure and shutting down zygotic induction after successful plasmid establishment.

### Delayed leading-region conversion enhances plasmid establishment in non-isogenic hosts

To determine whether modulation of early gene expression impacts plasmid establishment, we measured transfer efficiencies of F plasmid variants with altered leading-region conversion dynamics. Deletion or replacement of *ssp2* had no detectable effect on transfer efficiency between isogenic *E. coli* K12 donor and recipient strains, consistent with the absence of strong establishment barriers in this genetic background (Figure 4b).

Because leading-region genes are proposed to neutralize host defence systems encountered upon entry into new host strains, we next assessed plasmid transfer efficiency in non-isogenic recipient backgrounds. In *E. coli* strains HS, IAI1, B, and ED1a, plasmids lacking *ssp2* exhibited significantly reduced transfer efficiencies compared to the wild-type F plasmid, with strain-specific decreases ranging from 21-fold (HS & B) to undetectable transconjugant levels (ED1a) (Figure 4b). Consistently, the *ssp2* promoter controls the expression of *yfgB*, whose product shares 93% identity with the ArdB protein from plasmid R64, previously shown to function as an inhibitor of RM systems (Balabanov et al., 2012) which would not be produced in a Δ*ssp2* plasmid. To abolish the roadblock activity without impairing transcription of downstream genes, we replaced *ssp2* with *ssp3*, a ssDNA promoter of equivalent strength (Wen et al., 2025) but with no roadblock activity (Figure 4b). Replacement of *ssp2* with *ssp3* partially alleviated the loss of F transfer efficiencies for strains IAI1 and ED1a seen for a Δ*ssp2* plasmid. However, this effect was not observed in strains HS and B, suggesting that loss of roadblock activity at *ssp2* has a greater impact on plasmid transfer in these strains (Figure 4b).

These results indicate that delayed leading-region conversion and enhanced early gene expression provide a selective advantage during plasmid establishment in genetically diverse hosts. Notably, these recipient strains are known or predicted to encode distinct and heterogeneously distributed defence systems, including restriction-modification systems and other anti-plasmid barriers. Defence system prediction in these strains revealed that these four strains encode a variety of RM systems or related systems (i.e. BREX), prompting us to test the effect of individual systems on plasmid transfer. Transfer of a Δ*ssp2* plasmid in a strain encoding EcoGIII, a type II RM system (Fang et al., 2012), showed a 36-fold decreased transfer efficiency relative to a *wt* plasmid (Figure 4c). However, transfer of a *ssp2*::*ssp3* plasmid was similar to that of a *wt* plasmid, suggesting that the roadblock activity at *ssp2* does not affect plasmid evasion toward type II RM systems. On the other hand, transfer of a Δ*ssp2* plasmid in a strain encoding EcoKI, a type I RM, showed a 5-fold decreased transfer efficiency, which was not compensated when *ssp2* was replaced by *ssp3*, showing that the roadblock at *ssp2* provides an advantage for plasmid establishment under challenge by type I RM systems, likely by modulating the expression of the *yfgB* anti-RM gene and other potentially important leading genes located downstream of *ssp2* (Figure 4d).

## DISCUSSION

Successful dissemination of conjugative plasmids requires not only efficient DNA transfer from donor cells but also rapid and effective establishment in recipient cells. Here, we show that plasmids actively coordinate DNA processing and early gene expression to secure establishment in new host cells. By resolving the spatiotemporal dynamics of plasmid DNA conversion at single-cell resolution, we demonstrate that complementary strand synthesis is selectively delayed within the leading region, thereby extending zygotic induction and amplifying early plasmid gene expression. This delay is imposed by a single-stranded DNA promoter that acts as a transient replication roadblock and is resolved through the activity of the host helicase UvrD, revealing that plasmid establishment depends on a finely tuned interplay between plasmid-encoded regulatory elements and host DNA metabolism.

Our single-cell measurements refine the program of complementary strand synthesis that accompanies plasmid entry into recipient cells. Outside the leading region, ssDNA-to-dsDNA conversion scales linearly with distance from *oriT*, indicating a highly ordered conversion process. Because the transferred strand enters the recipient cell 5′ end first, this pattern is most consistent with a discontinuous mode of synthesis in which the incoming plasmid strand is repeatedly primed for DNA polymerase-mediated extension rather than converted from a single origin. These dynamics closely match the model proposed by Willetts and Wilkins (Willetts and Wilkins, 1984), in which the incoming ssDNA is rapidly coated by single-strand binding proteins and converted through successive priming events as it enters the cell.

Our rifampicin and UvrD experiments further clarify how host factors coordinate DNA processing with early gene expression. Inhibition of transcription did not block complementary strand synthesis, confirming that plasmid conversion relies exclusively on host replication machinery rather than *de novo* synthesis of plasmid-encoded proteins, as originally proposed by Hiraga (Hiraga and Saitoh, 1975). Moreover, rifampicin did not suppress the conversion delay in the leading region, ruling out the implication of *de novo* synthesized plasmid factors as well transcription-replication conflicts as its cause. Instead, the selective exacerbation of downstream conversion delays indicates that transcription at stem-loop-forming ssDNA promoters positively contributes to complementary strand synthesis, likely by generating RNA-DNA hybrids that serve as priming intermediates, as previously shown (Masai and Arai, 1997). In this context, delayed conversion arises from a structural barrier that must be actively resolved rather than from transcriptional interference. Consistent with this interpretation, we identify UvrD as a key host factor that enables replisome progression through the *ssp2*-dependent roadblock, revealing that plasmids exploit host DNA unwinding activities to overcome self-imposed obstacles. More broadly, variation in host DNA processing pathways, together with differences in leading-region regulatory architectures and gene repertoires, particularly anti-defence systems, is likely to be a major determinant of plasmid establishment efficiency, host range, and antibiotic resistance dissemination.

Delaying DNA conversion within the leading region provides an elegant solution to a central challenge of horizontal gene transfer: enabling the immediate deployment of protective functions upon entry while ensuring their rapid shutdown once establishment is secured. By prolonging the single-stranded state of the leading region, plasmids maximize early protective gene expression, a strategy that is particularly critical in non-isogenic hosts encoding diverse defence systems. Automatic shutdown of transcription upon conversion to double-stranded DNA ensures that this response is both potent and self-limiting, thereby minimizing fitness costs associated with sustained expression. Together, these findings reveal that complementary strand synthesis and early gene expression are tightly integrated into a unified regulatory strategy, allowing plasmids to actively tune early post-entry events. This coordination reframes plasmid transfer as a temporally regulated process in which precise control of timing, rather than transfer efficiency alone, emerges as a key determinant of establishment success and dissemination potential.

Several limitations should be acknowledged. Our analyses focus on the F plasmid, and the extent to which delayed DNA conversion is conserved across plasmid families remains unknown. In addition, although we identify UvrD as a key host factor, other replication proteins likely contribute to conversion initiation and progression and were not systematically explored. Finally, while delayed conversion enhances establishment in non-isogenic hosts, the specific roles of individual leading-region genes in counteracting distinct host defence pathways remain to be defined.

Beyond plasmid biology, our findings highlight single-stranded DNA promoters as a versatile regulatory module. By directly linking transcription to DNA structural state, these promoters function as intrinsic genetic timers that drive rapid, transient gene expression and self-silence upon double-strand synthesis, suggesting potential applications for engineering self-limiting genetic programs in synthetic biology.

## METHODS

### Strains, plasmids & growth conditions

Strains used in experiments are derived for *E. coli* MG1655 and listed in Table 1. Plasmids are listed in Table 2. Modifications of F plasmids were performed by recombineering in lambda red-competent strain DY330 (Yu et al., 2000), followed by mating transfer in MG1655 and cassette excision with Flp-encoding plasmid pCP20 (Datsenko and Wanner, 2000). The Δ*uvrD* allele was transduced from KEIO strain JW3786 (Baba et al., 2006) using phage P1vir. Scarless deletion of *ssp2* and its replacement with *ssp3* was performed by double recombination using the SacB-encoding plasmid pKanSac. Cells were grown in Lysogeny Broth (LB) or Agar (LA) according to Lennox’s formula (5 g/l yeast extract, 10 g/l tryptone, 5 g/l sodium chloride, and 15 g/l agar for solid medium). For microscopy experiments, cells were grown in M9 medium (7 g/l disodium phosphate heptahydrate, 3 g/l monopotassium phosphate, 1 g/l ammonium chloride, 0.5 g/l sodium chloride, 1 mM magnesium sulfate & 80 µM calcium chloride) supplemented with 2 g/l glucose, 4 g/l casamino acids and 400 µg/l thiamine hydrocholoride ; this medium is referred to as M9GC. When appropriate, antibiotics were used at the following concentrations: carbenicillin disodium 25 µg/ml, chloramphenicol 20 µg/ml, kanamycin sulfate 50 µg/ml, streptomycin 20µg/ml & tetracycline hydrochloride 10 µg/ml.

**Table 1.**
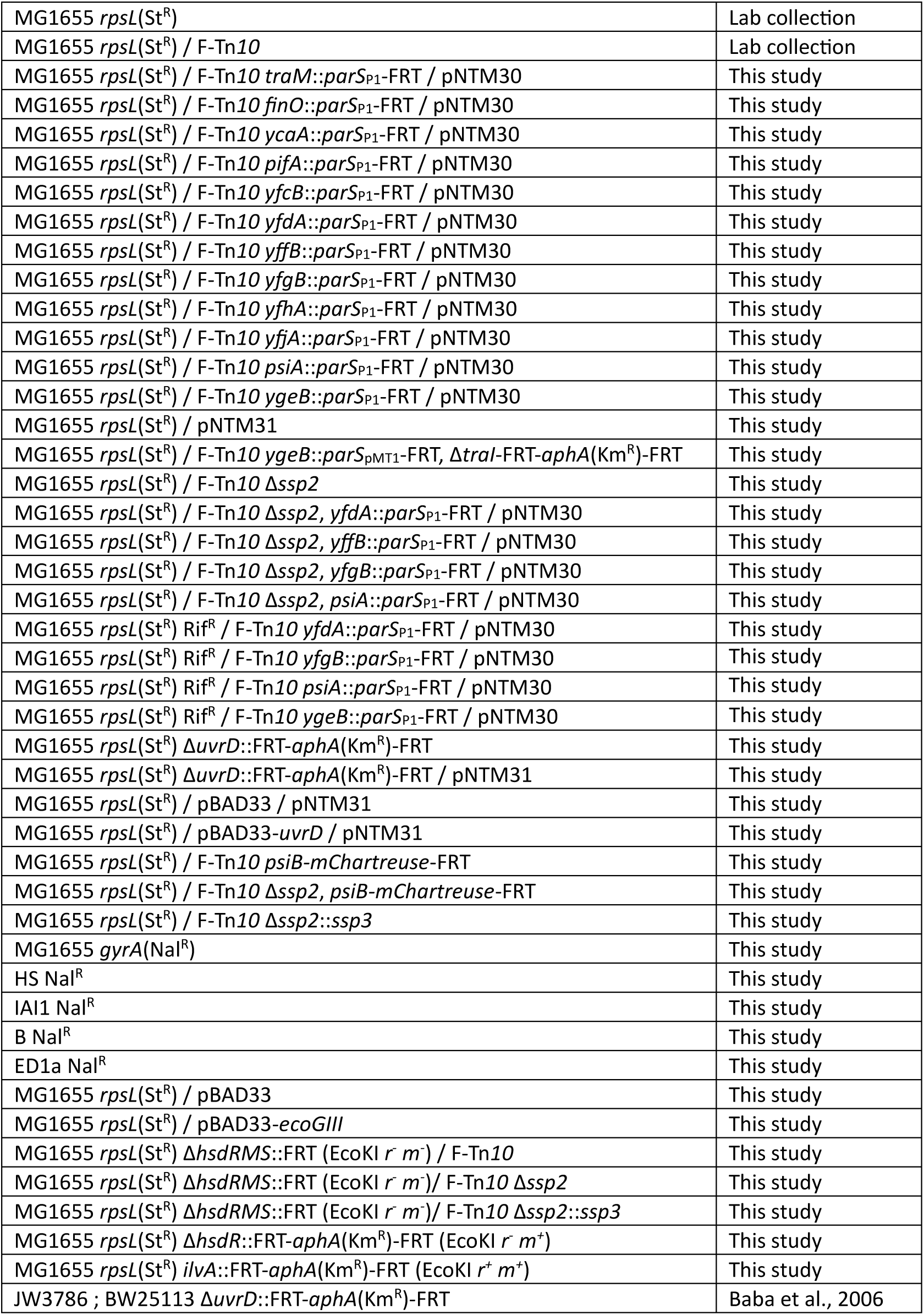
Strains used in this study

**Table 2.** Plasmid vectors used in this study

|  |  |  |
| --- | --- | --- |
| pNTM30 | *ori<sub>pSC101</sub> bla(Amp<sup>R</sup>) ssb-mLychee* | This study |
| pNTM31 | *ori<sub>pSC101</sub> bla(Amp<sup>R</sup>) ssb-mLychee parB<sub>P1</sub>-mChartreuse* | This study |
| pBAD33 | *ori<sub>p15a</sub> cat(Cml<sup>R</sup>) ParaBAD* | Lab collection |
| pBAD33-uvrD | *ori<sub>p15a</sub> cat(Cml<sup>R</sup>) ParaBAD-uvrD* | This study |
| pBAD33-ecoGIII | *ori<sub>p15a</sub> cat(Cml<sup>R</sup>) ParaBAD-ecoGIII(*r<sup>+</sup> m<sup>+</sup>*)* | Fang et al., 2012 |
| pKanSac | *ori<sub>R6K</sub> aphA(Kan<sup>R</sup>) sacB(Suc<sup>S</sup>)* | This study |

### Microscopic analysis of conversion delay

Donors and recipient cultures were grown in M9GC to OD 0.7-0.9, mixed with a 1:1 ratio and spotted on M9GC pads gelled with 1% agarose. Pads were sealed between microscopy slides and coverslips with 125 µl Gene Frames (Thermo Scientific) and imaged at 37°C with a Eclipse Ti2 microscope (Nikon) equipped with a Plan Apo λ 100×/1.45 objective, a motorized stage with Z-drift correction (Perfect Focus System, Nikon), heating (Okolab), a LED light source (Spectra X, Lumencor), and a sCMOS camera (Orca-Fusion BT, Hamamatsu). Images were acquired every 30s using GFP (474/27 excitation filter, 515/30 emission filter, 100 ms exposure 50% power) and RFP channels (553/24 excitation filter, 580LP emission filter, 100 ms exposure, 12% power). Conversion delay was calculated as the time delay between the appearance of Ssb-mLychee foci and that of ParB-mChartreuse foci in individual mating events.

### DNA 3’ end capture

Short duration, high-volume (30 min, 20 ml) 1:1 mating reactions were quenched on ice-water slurry, centrifuged (5 min, 5000 RCF) and stored at -20°C for future use. Plasmid DNA was extracted from pellets using standard miniprep procedures (NucleoSpin Plasmid, Macherey-Nagel). A 20-ng aliquot of DNA in TdT buffer (New England Biolabs) supplemented with 250 µm CoCl2 and 200 µM dCTP was denatured (98°C, 5min) and chilled on ice-water slurry. Poly-dC tailing was performed by the addition of 20 U TdT at 37°C for 30 min, followed by TdT inactivation (70°C, 10 min). A 50 µl PCR reaction was setup using a ssp2-specific primer (GTAAAACGACGGCCGATTGTATTGAACATATCCTGTC, 200 nM), an anchor primer containing a 3’ dGI tract (CAGGAAACAGCTATGACGGGIIGGGIIGGGIIG, 400 nM), 2 µl of tailing reaction and 0.125 U DreamTaq (Thermo Scientific). A second nested PCR reaction was performed with 2 µl of PCR product using primers yffB up F (CGATGAGGACGACGAAAACC, 200 nM) and M13 R (CAGGAAACAGCTATGAC, 400 nM). Reaction products were separated by electrophoresis on 1.2% agarose 0.5x TAE gels, excised from the gels, purified (NucleoSpin PCR, Macherey-Nagel) and sequenced (Sanger Sequencing, Microsynth).

### Conjugation efficiency assays

Overnight cultures grown in LB were centrifuged and resuspended in fresh LB medium to an OD600 of 2.0. Donor and recipient cultures were mixed in a 1:1 ratio and 100 µl were spotted on 0.45 µm cellulose ester filters (HAWP02500, Millipore) adsorbed on LA plates. After a 120 min incubation at 37°C, matings were resuspended in 1 ml LB, serially diluted in LB and spotted on LA plates containing appropriate antibiotics to select recipients or transconjugants. Recipient colonies were then patched on LA plates containing tetracycline to select for transconjugants. Frequencies of transconjugants were calculated as the number of transconjugant patched divided by the number of patched recipient colonies.

### Data availability

All data to understand and assess the conclusions of this research are available in the main text and figures.

## Acknowledgements

We thank members from the Lesterlin & Yamaichi team for valuable input during the project progress. Strain JW3786 was sourced from the KEIO collection, kindly provided by the National BioResource Project, National Institute of Genetics, Japan.

## Author contribution statement

C.L., N.F. and Y.Y. conceptualized the study. N.F. and C.L. analysed the data, wrote the paper and prepared the figures. N.F conducted experiments. C.L. and Y.Y. provided funding.

